# Cellular Mechanisms Underlying Embryonic Retinal Waves

**DOI:** 10.1101/2022.08.14.503889

**Authors:** Christiane Voufo, Andy Quaen Chen, Benjamin E. Smith, Marla B. Feller, Alexandre Tiriac

## Abstract

Spontaneous activity is a hallmark of developing neural systems. In the retina, spontaneous activity comes in the form of retinal waves, comprised of three stages persisting from embryonic day 16 (E16) to eye opening at postnatal day 14 (P14). Though postnatal retinal waves have been well characterized, little is known about the spatiotemporal properties or the mechanisms mediating embryonic retinal waves, designated Stage 1 waves. Using a custom-built macroscope to record spontaneous calcium transients from whole embryonic retinas, we show that Stage 1 waves are initiated at several locations across the retina and propagate across finite regions of a broad range of areas. A gap junction antagonist, meclofenamic acid, reduced the frequency and size of Stage 1 waves but did not abolish them. The general nAChR antagonist, hexamethonium blocked Stage 1 waves, while they persisted in the presence of α4β2 nAChR antagonist dihydro-ß-erythroidine, indicating that the spatiotemporal properties of Stage 1 waves are mediated by a complex circuitry involving subtypes of nAChRs and gap junctions. Stage 1 waves in mice lacking the β2 subunit of the nAChRs (β2-nAChR-KO) were reduced, but in contrast to WT mice, they persisted in the hexamethonium and were completely blocked by meclofenamic acid. To assay the impact of Stage 1 waves on retinal development, we compared the spatial distribution of a subtype of retinal ganglion cells, intrinsically photosensitive retinal ganglion cells (ipRGCs) in WT and β2-nAChR-KO mice. We found that the developmental decrease of ipRGC density is preserved between WT and β2-nAChR-KO mice, indicating that processes regulating ipRGC distribution are not influenced by spontaneous activity.

## INTRODUCTION

Throughout the developing nervous system, spontaneous activity is observed before neural circuits are fully formed and sensory transduction begins (Blankenship and Feller, 2010; Luhmann and Khazipov, 2018; Akin and Zipursky, 2020; Martini et al., 2021). This activity is implicated in several development events, including cell death, maturation of functional circuits, and refinement of projection neurons in their targets in the brain (Kirkby et al., 2013; Blanquie et al., 2017a; Fujimoto et al., 2019). This is well studied in the developing visual system, where prior to the maturation of vison, laterally propagating spontaneous depolarizations sweep across retinal ganglion cells (RGCs), a pattern referred to as retinal waves (Wong, 1999). Retinal waves drive eye specific segregation and retinotopic refinement of retinal projections to the dorsal lateral geniculate nucleus and superior colliculus (Ackman and Crair, 2014; Arroyo and Feller, 2016). Retinal waves also play a role in the maturation of direction selective circuits within the retina itself (Tiriac et al., 2022) as well as the development of retinal vasculature (Weiner et al., 2019; Biswas et al., 2020).

Retinal waves are present throughout mouse retinal development, starting as early as embryonic day 16 and persisting until postnatal day 14, which is around the time of eye opening (Blankenship and Feller, 2010; Feller and Kerschensteiner, 2013; Choi et al., 2021). As the retina develops, the circuits that mediate waves change. Stage 2 retinal waves, observed between postnatal day 1 and 10 (P1-10), are mediated via activation of nicotinic acetylcholine receptors (nAChRs) by acetylcholine (ACh) released from starburst amacrine cells (SACs). Stage 2 waves are propagated through a network of SACs that themselves express nAChRs during development (Zheng et al., 2004). Stage 3 retinal waves, observed between P10-14, are mediated via activation of ionotropic glutamate receptors by glutamate released from bipolar cells. Stage 3 waves are propagated through a network of bipolar and amacrine cells via glutamate transmission and gap junction coupling. Previous findings show that Stage 3 wave frequency is enhanced by light stimulation, indicating that the photoreceptor-depolarizations are incorporated into Stage 3 wave-generating circuits (Tiriac et al., 2018).

Retinal waves are also present in embryonic retinas. In mice, Stage 1 waves are observed between embryonic day 16 and 18 (E16-18) (Bansal et al., 2000) while in rabbit they are present at E22 (Syed et al., 2004). In rabbit, Stage 1 waves persist in the presence of pharmacological antagonists of fast neurotransmitters and are blocked by gap junction antagonists (Syed et al., 2004). In mice, Stage 1 waves consist of large propagating waves and small non-propagating events (Bansal et al., 2000). The application of nAChR antagonist inhibits larger propagating events (Bansal et al., 2000). Though there is no anatomical evidence of synapses as early as E16-18, recent work has shown that SACs are present embryonically, begin to migrate to the inner nuclear layer (INL), and send projections to the inner plexiform layer (IPL) guided by homotypic contacts (Ray et al., 2018). Hence cholinergic signaling is likely occurring via the volumetric release of ACh. Exactly how gap junctions and cholinergic signaling set the spatiotemporal properties of Stage 1 waves remains to be understood.

Embryonic retinal development is a dynamic process involving cell death, proliferation, specification, migration, and process outgrowth. Specifically, concurrent with Stage 1 waves are RGC death and RGC subtype specific reduction in proliferation (Marcucci et al., 2019; Shekhar et al., 2022). The latter has been well characterized in intrinsically photosensitive retinal ganglion cells (ipRGCs) (McNeill et al., 2011) and the former has been observed across the general RGC population (Bähr, 2000; Braunger et al., 2014). Whether these processes are influenced by activity remain unknown.

Here we describe the spatiotemporal properties of Stage 1 waves across the whole retina using a novel macroscope. In addition, we report a role for nAChR signaling and gap junction coupling in setting up these properties, which differ from Stage 2 waves. The β2-nAChR-KO mice, the canonical mouse model for the studying the role of Stage 2 waves in developmental processes also exhibits reduced wave activity. Finally, we use β2-nAChR-KO mice to demonstrate that the regulation of ipRGC number is a wave-independent process.

## RESULTS

### Macroscope imaging reveals that Stage 1 retinal waves have complex spatiotemporal properties

The mouse retina at E16-18 exhibits spontaneous correlated events (Bansal et al., 2000; Syed at al., 2004), termed Stage 1 retinal waves, despite the immature state of retinal circuits. At E16-18, several postmitotic cell types are present in the retina, including broad classes of retinal ganglion cells (RGCs) and amacrine cells (ACs), as well as proliferating progenitors which will go on to produce cells such as rods, bipolar cells, and Muller glia postnatally (Fig 1A)(Cepko, 2014). By E17, migrating SACs release ACh (Wong, 1995) and they, along with RGCs, express nicotinic acetylcholine receptors; however, there are no chemical synaptic structures (Hoon et al., 2014). There is extensive gap junction coupling between RGCs as well as between progenitor cells (reviewed in Cook and Bekker, Physiology, 2009), which have been proposed to be the primary substrate mediating Stage 1 waves (Catsicas et al., 1998; Bansal et al., 2000; Syed et al., 2004).

**Figure 1.**
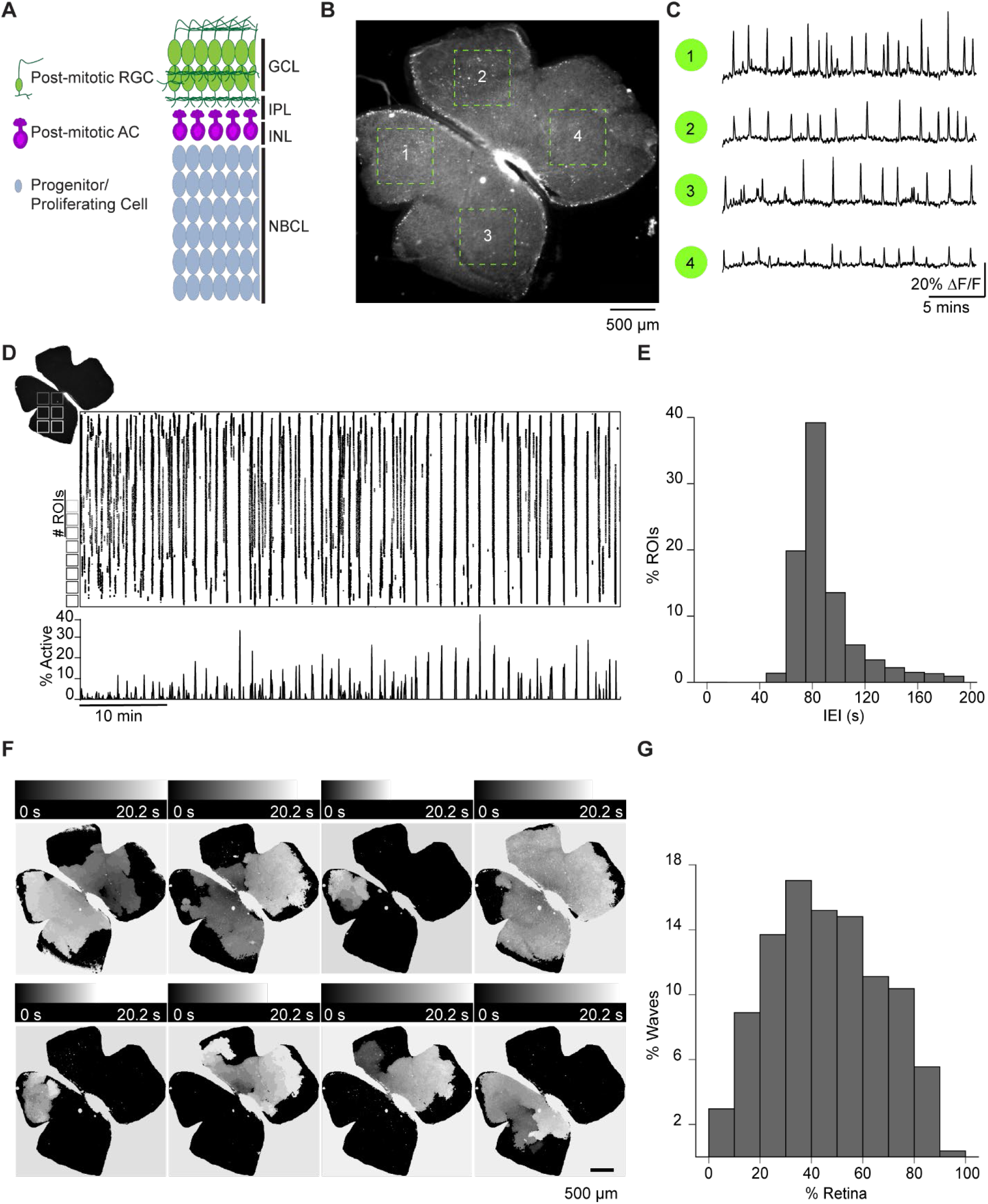
Spatiotemporal characteristics of embryonic waves. **A**, schematized cross section of an E16 retina when Stage 1 waves begin. Green cells represent post-mitotic RGCs. Magenta cells represent post-mitotic ACs. Gray cells, represent progenitor cells and proliferating cells. GCL, ganglion cell layer; IPL, inner plexiform layer; INL, inner nuclear layer; NBCL, neuroblastic cell layer. **B**, Macroscope image of the baseline fluorescence of an E17 bath loaded with Cal 520 (FOV 4.7 mm x 4.7 mm). Dashed green squares and numbers indicate the four example ROIs (600 µm x 600 µm represented in **C. C**, 20 min time course of the normalized change in fluorescence (ΔF/F) observed in the four ROIs indicated in **B. D**, top left: outline of an example retina. For subsequent analysis, the retina was divided into 10 µm x 10 µm (not drawn to scale) squares ranging from 8,179 - 12,692 squares, depending on the size of the retina. Middle: raster plot of calcium transients > 30% ΔF/F for bath loaded retinas and 50% for GCaMP6s expressing retinas. Bottom: Area plot summarizing the percentage of ROIs active throughout the time course of the recording (1 hour). **E**, normalized histogram showing the distribution of the percent of ROIs with inter-event-intervals (IEIs) ranging from 40 - 200s. **F**, heatmap showing the temporal progression and spread of 8 example events observed with epifluorescent calcium imaging on the macroscope. Scale depicts timescale of propagation, black = start of propagation and white = end of propagation. **G**, normalized histogram showing the distribution of events with an area ranging from 0-100% of the retina.

To understand how cholinergic signaling and gap junction coupling instruct the spatiotemporal properties of Stage 1 waves, we first characterize these waves via the following properties: frequency, area, speed, and the distribution of initiation sites. Stage 1 retinal waves were recorded using a custom built epifluorescent macroscope. The macroscope had a large FOV of 4.7 mm x 4.7 mm (22.09 mm^2^) while maintaining subcellular resolution (3.0 µm), which allowed for recording of activity across the entire retina (Figure 1B & C). Retinas were isolated from E16-18 mice that were either bath loaded with the organic calcium dye Cal 520, or from mice expressing the genetically encoded calcium indicator GCaMP6s under the *Vglut2* promoter (Vglut2::GCaMP6s). The earliest age at which we could detect reliable wave-like events was E16 (Supp Movie 1).

To assess the spatiotemporal properties of Stage 1 waves, we divided the retinal surface into small square ROIs (roughly 10 µm x10 µm), about 7 µm apart. ΔF/F traces for each ROI were rasterized based on an event detection algorithm which identified the timing of peak changes in fluorescence (Figure 1D). These rasterized events were used for subsequent analysis. We first computed the time between spontaneous events by measuring the inter-event interval (IEI) for each ROI and found that the distribution peaked at around 80 seconds (Figure 1E). We also described the size of individual retinal waves by computing the percent of ROIs that participated in each wave. The distribution of wave sizes was broad, ranging from 6% to 90% of the retina, with a mean and standard deviation of 46 ± 21% (Figure 1F & G).

In addition to propagating events, we also observed small non-propagating events (defined as events that covered less than 4 ROIs) that exhibited correlated spontaneous increases in [Ca^2+^] (Supp Movie 2). These events were like those described previously in (Bansal et al., 2000). Note, these small non-propagating events were included in the IEI but not in the analysis of propagating wave sizes.

Finally, we characterized the average propagation speed of Stage 1 waves and found that large propagating waves had a propagation speed ranging from 145 μm/s to 237 μm/s (Average ± SD: 181 ± 24 μm/s) (Figure 3C), similar to Stage 2 waves we observed with an average speed of 177 μm/s ± 62 μm/s (see Table 1 for wave speed summary data & Figure S1 for a description of speed calculation). We also measured the distribution of Stage 1 wave initiation sites in WT retinas and saw no evidence of an initiation site bias (Figure S2).

**Table 1:**
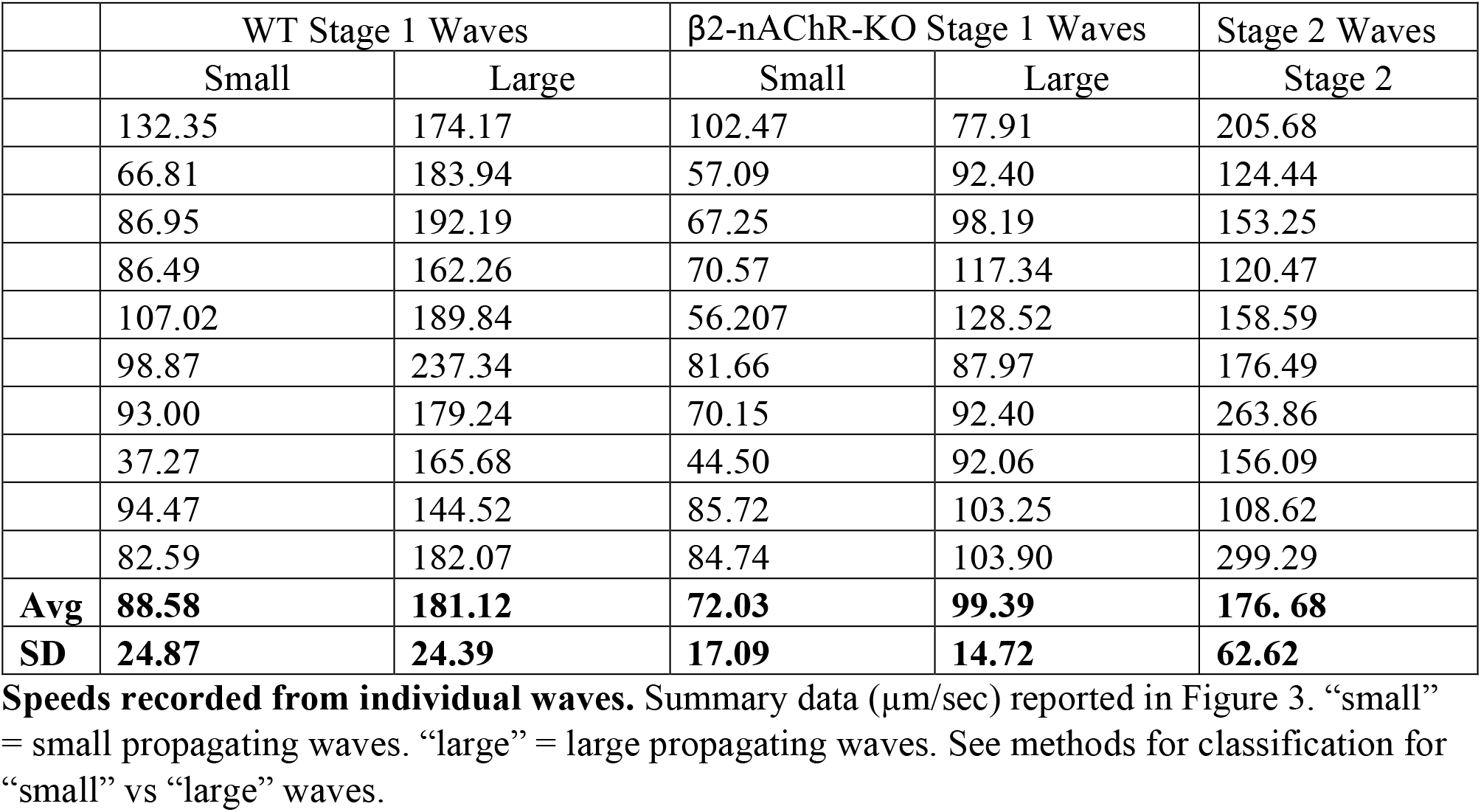
Stage 1 small and large wave speed in WT and β2-nAChR-KO, plus Stage 2 wave speed.

| | WT Stage 1 Waves | | $\beta 2$-nAChR-KO Stage 1 Waves | | Stage 2 Waves |
| --- | --- | --- | --- | --- | --- |
|  | Small | Large | Small | Large | Stage 2 |
|  | 132.35 | 174.17 | 102.47 | 77.91 | 205.68 |
|  | 66.81 | 183.94 | 57.09 | 92.40 | 124.44 |
|  | 86.95 | 192.19 | 67.25 | 98.19 | 153.25 |
|  | 86.49 | 162.26 | 70.57 | 117.34 | 120.47 |
|  | 107.02 | 189.84 | 56.207 | 128.52 | 158.59 |
|  | 98.87 | 237.34 | 81.66 | 87.97 | 176.49 |
|  | 93.00 | 179.24 | 70.15 | 92.40 | 263.86 |
|  | 37.27 | 165.68 | 44.50 | 92.06 | 156.09 |
|  | 94.47 | 144.52 | 85.72 | 103.25 | 108.62 |
|  | 82.59 | 182.07 | 84.74 | 103.90 | 299.29 |
| **Avg** | **88.58** | **181.12** | **72.03** | **99.39** | **176.68** |
| **SD** | **24.87** | **24.39** | **17.09** | **14.72** | **62.62** |

### nAChRs and gap junctions are important for setting the frequency and area of Stage 1 waves

Previous work done in mice has shown that large propagating waves observed in the embryonic retina are sensitive to curare, a competitive antagonist for nAChR (Bansal, 2000). However, in rabbit, blockade of all fast neurotransmitter receptor, including nAChRs, had no impact on Stage 1 wave frequency (Syed et al, 2004). Rather, wave frequency was blocked after the application of 18β-glycyrrhetinic acid, a gap junction antagonist (Syed et al., 2004). To determine the relative role of nAChRs and gap junctions on the frequency and area of Stage 1 waves in mice, we used two-photon calcium imaging and pharmacology in retinas isolated from E16-18 mouse pups bath loaded with Cal 520. We found that the frequency and area of Stage 1 waves was significantly reduced in the presence of gap junction blocker meclofenamic acid (MFA, 50μM), with a corresponding reduction in cell participation (Fig 2A & 2F). We also found a small but significant reduction in wave amplitude, quantified as the average maximum response amplitude of all cells participating in individual waves (Fig 2H). In addition, Stage 1 waves were blocked in the presence of hexamethonium (Fig 2B; Hex, 100μM), a non-selective nAChR antagonist, as well as in epibatidine (Fig 2C; EPB, 10 nM), a nAChR agonist that potently desensitizes all nAChRs (Spang et al., 2000; Corrie et al., 2020).

**Figure 2.**
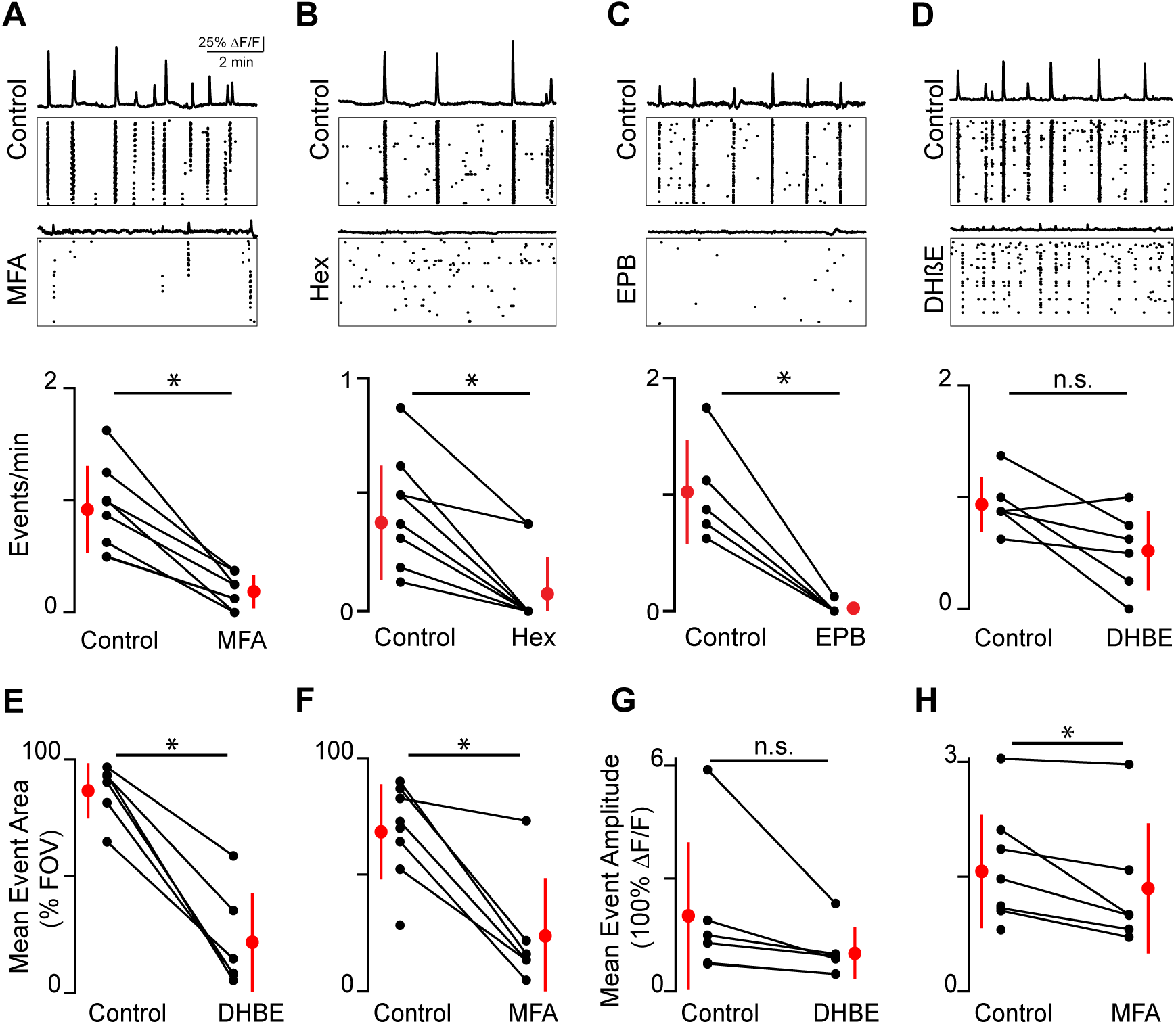
Embryonic waves are mediated by gap junction and cholinergic circuits. **A**, top: ΔF/F time course of events observed in the field of view (FOV) and raster plot of neuronal calcium transients > 15% ΔF/F in control. Middle: ΔF/F time course of events observed in the FOV and raster plot of neuronal calcium transients > 15% ΔF/F in MFA (50 µM). Bottom: Summary plot showing frequency of events in control and MFA. Red dots and lines = mean and standard deviation, respectively. Asterisks represents significant effects. **B**, top: ΔF/F time course of events observed in the FOV and raster plot of neuronal calcium transients > 15% ΔF/F in control. middle: ΔF/F time course of events observed in the FOV and raster plot of neuronal calcium transients > 15% ΔF/F in Hex (100 µM). Bottom: Summary plot showing frequency of events in control and Hex. n = 11 retinas (7 mice); p = 5.33e^-4^. Paired t-test. **C**, top: ΔF/F time course of events observed in the FOV and raster plot of neuronal calcium transients > 15% ΔF/F in control. Middle: ΔF/F time course of events observed in the FOV and raster plot of neuronal calcium transients > 15% ΔF/F in EPB (10 nM). Bottom: Summary plot showing frequency of events in control and EPB. n = 5 retinas (5 mice); p = 4.8e^-3^. Paired t-test. **D**, top: ΔF/F time course of events observed in the FOV and raster plot of neuronal calcium transients > 15% ΔF/F in control. Middle: ΔF/F time course of events observed in the FOV and raster plot of neuronal calcium transients > 15% ΔF/F in DHβE (8 μM). Bottom: Summary plot showing frequency of events in control and DHβE. **E**, mean event area in control and DHβE. **F**, mean event area in control and MFA. Paired t-test (**D-F**). **G**, mean event amplitude in control and DHβE. **H**, mean event amplitude in control and MFA. n = 6 retinas (6 mice) (**D, E** & **G**); p = 0.05 (**D**); p = 4.58e^-4^ (**E**); p = 0.12 (**G**). n = 8 retinas (6 mice) (**A, F** & **H**); p = 4.81e^-4^(**A**); p = 0.03 (**G**); p = 0.02 (**H**). Paired t-test.

To further explore the role of different subtypes of nAChRs on the frequency and area of Stage 1 waves, we bath applied dihydro-ß-erythroidine hydrobromide (DHβE, 8μM), which preferentially targets nAChRs containing α4 and β2-subunits (Harvey and Luetje, 1996; Harvey et al., 1996) and is a potent blocker of Stage 2 retinal waves (Ford et al., 2012). We found that DHβE did not prevent the generation of Stage 1 retinal waves (Figure 2D). Waves that persisted in the presence of DHβE were smaller and had comparable amplitudes to those observed in control (Figures 2E and 2G). This suggests that different subunits of nAChRs play distinct roles in the propagation of large events.

### Stage 1 waves persist in β2-nAChR knock-out mice but with reduced frequency and area

Our results thus far indicate that both the frequency and area of Stage 1 retinal waves are modulated by nAChR signaling comprised of different subunits as well as gap junction coupling. To further differentiate the role of different nAChRs, we characterized mice where the β2 subunit of the nicotinic acetylcholine receptor is genetically ablated (β2-nAChR-KO). β2-nAChR-KO mice have severely disrupted Stage 2 retinal waves (Bansal et al., 2000; Rossi et al., 2001) and have served as a model system for assessing the role of Stage 2 waves in driving different developmental events (Ackman and Crair, 2014; Burbridge et al., 2014).

We observed that β2-nAChR-KO retinas exhibited Stage 1 waves with different spatiotemporal properties than WT retinas (Figure 3A&B, see also Supp Movie 3). Specifically, β2-nAChR-KO retinas exhibited longer IEIs (Figure 3D&E), and individual waves propagated over smaller areas β2-nAChR-KO (Figure 3E). In sharp contrast to WT retina, waves in the β2-nAChR-KO retina were unaffected by the addition of Hex but showed a significant reduction in both area and frequency with the application of MFA (Figure 3D&E). These results suggest that in the absence of β2-nAChRs, gap junctions become a major component of the Stage 1 wave generation mechanism. Furthermore, we found that Stage 1 waves in the β2-nAChR-KO propagated at a significantly slower speed than those observed in WT mice, (Average ± SD: 99 ± 15 µm/sec compared to 181 ± 24 µm/sec for WT, n= 10 waves per condition, p = 3.9073e^-8^) (Figure 3C), consistent with a distinct mechanism of propagation.

**Figure 3.**
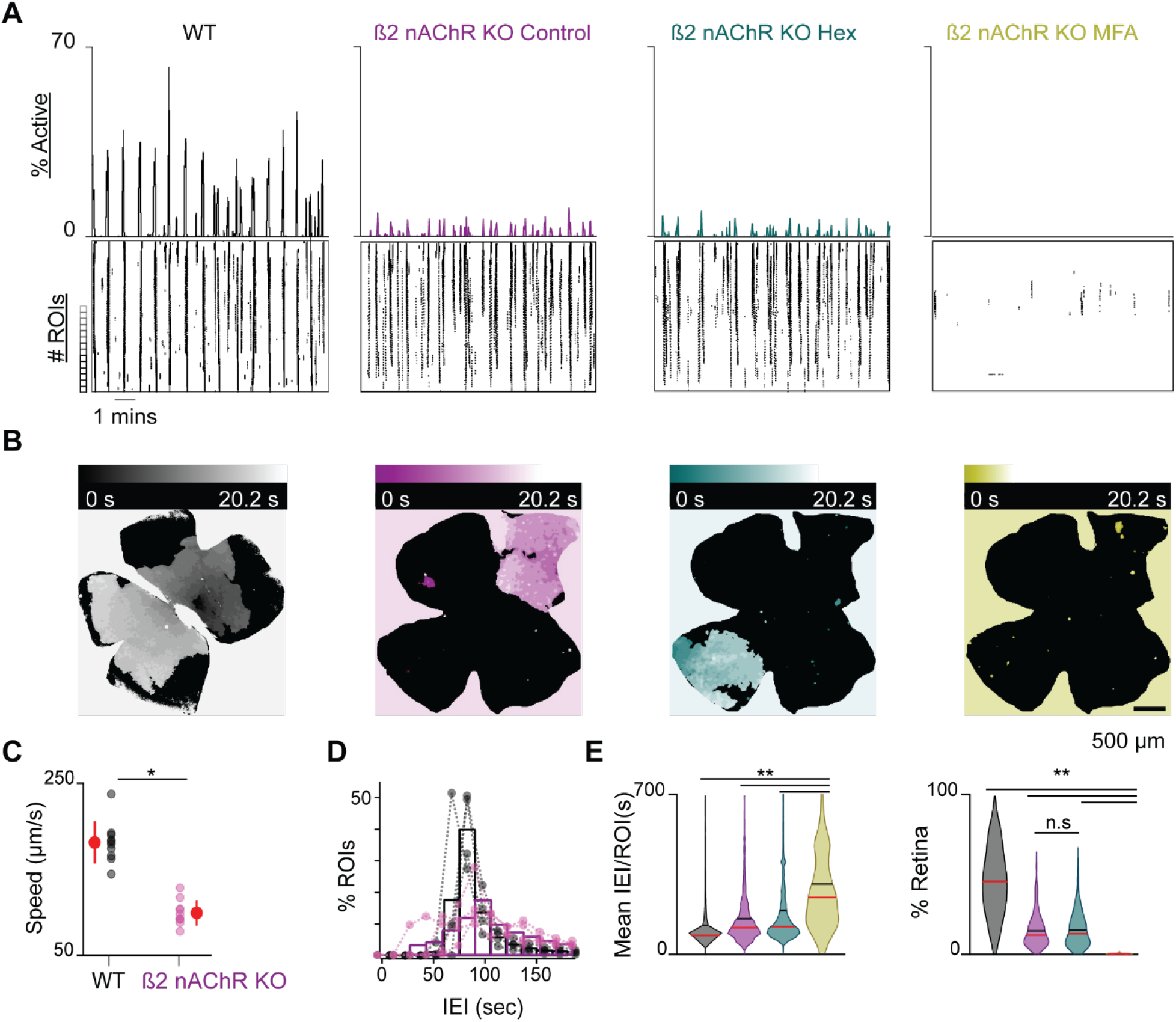
β2-nAChR-KO mice have reduced stage 1 wave activity. **A**, top, area plot summarizing the percentage of ROIs active throughout the time course of the recording; bottom, raster plot of calcium transients > 50% ΔF/F for GCaMP6s expressing retinas. Black: WT control; magenta: β2-nAChR-KO control; teal: β2-nAChR-KO in Hex; yellow: β2-nAChR-KO in MFA. **B**, heatmap showing the temporal progression of a propagating event observed using epifluorescent calcium imaging on the macroscope across experimental conditions. Scale depicts timescale of propagation, dark colors = start of propagation and white = end of propagation. **C**, summary plot of wave speed in WT and β2-nAChR-KO retinas in control conditions. n = 10 waves; p = 3.9e^-8^. Unpaired t-test. **D**, mean IEI/ ROI in WT and β2-nAChR-KO retinas in control conditions. Black bar = mean; red bar = median. **E**, violin plots summarizing the distribution of mean IEI/ ROI (right) and event area (left) across experimental conditions. n = 4 (WT), n = 5 (β2-nAChR-KO) (**D** & **E**); **p < 0.01. One-way ANOVA, followed by Tukey-Kramer post hoc test (right); Kruskal-Wallis, followed by Dunn-Sidak post hoc test (left).

Finally, we wanted to test whether the properties of Stage 1 waves in WT or β2-nAChR-KO were influenced by light activation of ipRGCs via the 474 nm imaging light used to excite the calcium dye on the macroscope. ipRGCs have been shown to be responsive to 476 nm light during both embryonic and postnatal development (Emanuel and Do, 2015; Verweij et al., 2019). Previous studies have shown a light dependent increase in the wave frequency of Stage 2 waves in the β2-nAChR-KO (Kirkby, 2013). This light modulation of the spatiotemporal properties of β2-nAChR-KO Stage 2 waves was shown to depend on ipRGC melanopsin expression, as well as an increase in gap junction conductance between ipRGCs and other RGCs (Kirkby and Feller, 2013; Arroyo et al., 2016). However, we found no difference in the frequency of waves recorded using a two-photon microscope (based on 920 nm illumination) to those recorded on the macroscope and in either the WT or β2 nAChR-KO retinas (Figure S3). Hence, we conclude that light activation of ipRGCs does not significantly influence the spatiotemporal properties of Stage 1 retinal waves in either WT or the β2 nAChR-KO retinas. These results are consistent with the fact that light stimulation of the retina does not modulate the frequency of Stage 2 waves and only begins to do so when conventional photoreceptors come online during Stage 3 waves (Tiriac et al, 2018).

### ipRGCs participate in Stage 1 waves but their number and distribution are not altered in the β2-nAChRs-KO

ipRGCs begin to proliferate from E10-E12 and show a decline in proliferation after E15 (Lucas, 2019; McNeill, 2012). They start expressing melanopsin at E15 and show responses to light as early as E17.5 (McNeill, 2012) (Verweij et al., 2019). ipRGCs, like all RGCs, also undergo a period of dramatic cell death during the first two postnatal weeks of development, the majority occurring during the first postnatal week (Braunger et al., 2014; Abed et al., 2022); however, the exact mechanism regulating the cell death is unknown. To test whether Stage 1 and Stage 2 retinal waves play a role in retinal development, we assessed the impact of disrupting waves on the number and distribution of ipRGCs.

We first set out to determine whether ipRGCs participate in Stage 1 waves. To do this, we conducted two-photon calcium imaging of RGCs in the ganglion cell layer (GCL) of retinas isolated from Opn4^Cre/+^; tdTomato^fl/fl^ E16-18 mice (Ecker et al., 2010), which express tdTomato in ipRGCs (Figure 4A) enabling us to assess the differential participation of RGCs and ipRGCs during Stage 1 waves (Figure 4B). On average, both RGCs and ipRGCs participated in most waves with no significant differences between the two groups (Figure 4C; Average ±SD: RGCs 78.78±21.48%; ipRGCs 84.64±16.86%). We also found no significant differences in the amplitude of the calcium response that RGCs and ipRGCs exhibit in response to Stage 1 waves (Figure 4D). Hence, ipRGCs are depolarized by Stage 1 waves similarly to Stage 2 waves (Chew et al., 2017; Caval-Holme and Feller, 2019; Caval-Holme et al., 2022).

**Figure 4.**
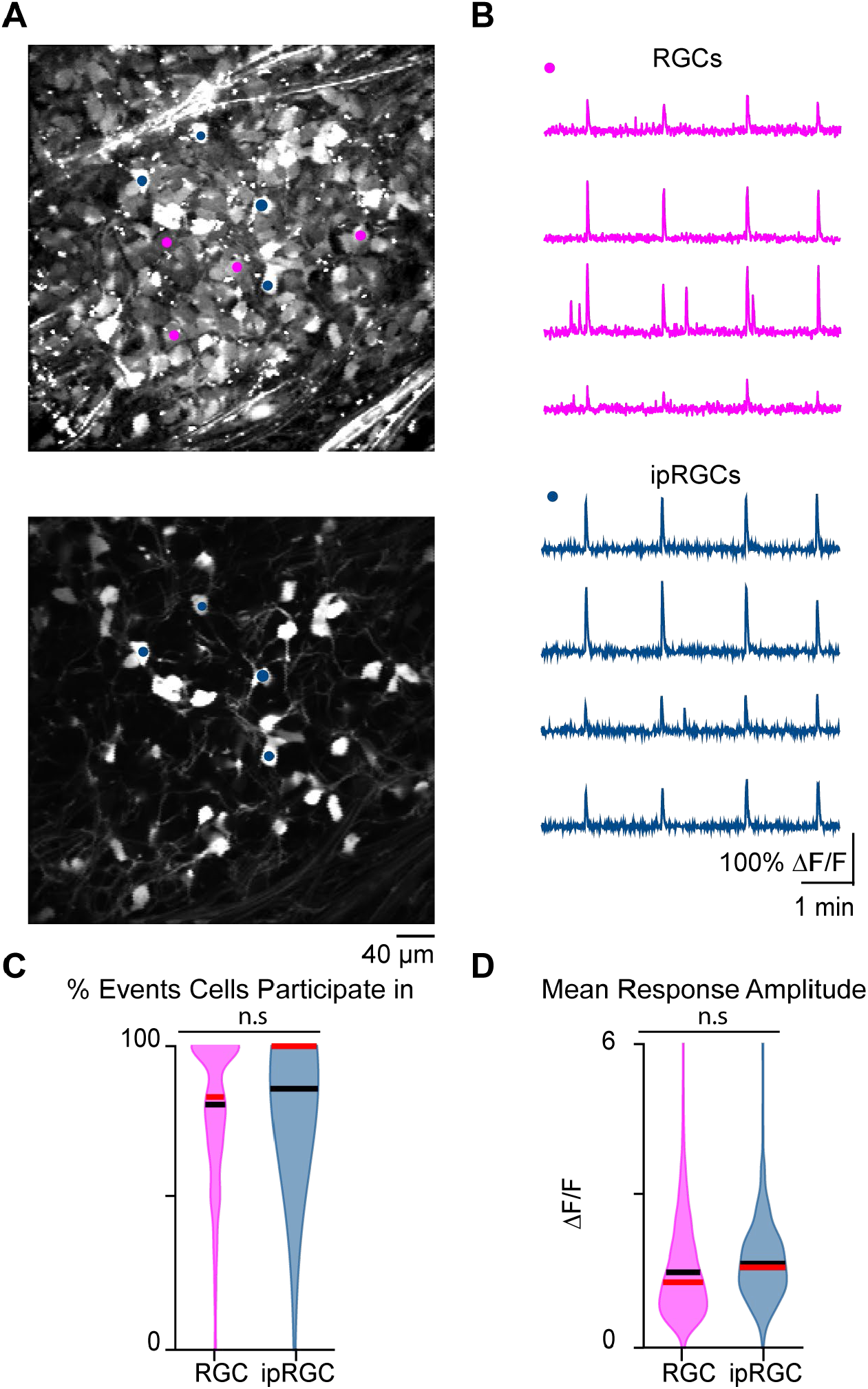
Stage 1 waves robustly recruit ipRGCs and the general RGC population. **A**. *Top:* Example FOV where pixel intensity was averaged across all frames (712) to get representative image of Cal 520 bath loaded RGCs. Pink dots correspond to four example RGCs and blue dots correspond to four example ipRGCs *Bottom:* same example FOV from the top image, with tdTom signal averaged across all frames. Blue dots the same as those in the top image. **B**. *Top:* traces of the four example RGCs marked by the pink circles in **A**. *Bottom:* traces of four example ipRGCs marked by the blue circles in **A. C**. Violin plot of percentage of waves each cell participated in. n = 20 retinas (13 mice); p = 0.09. Unpaired t- test. **D**. Violin plot of mean event amplitude/cell/FOV. n = 20 retinas (13 mice); p = 0.06. Unpaired t-test. Black bar = mean; red bar = median.

Since β2-nAChR-KO retinas exhibited reduced retinal activity during both embryonic and early postnatal development, we used this mouse as a model to determine whether normal Stage 1 and Stage 2 wave activity is important for regulating the number and distribution of ipRGCs. To this end, we isolated retinas from Opn4^Cre/+^;tdTomato^fl/fl^ and Opn4^Cre/+^;tdTomato^fl/fl^; (β2-nAChR^-/-^) at P1 and P7. These retinas express tdTomato in all melanopsin-expressing cells regardless of subtype. Retinas were imaged using wide-field epifluorescence microscopy (Figures 5A and B).

**Figure 5.**
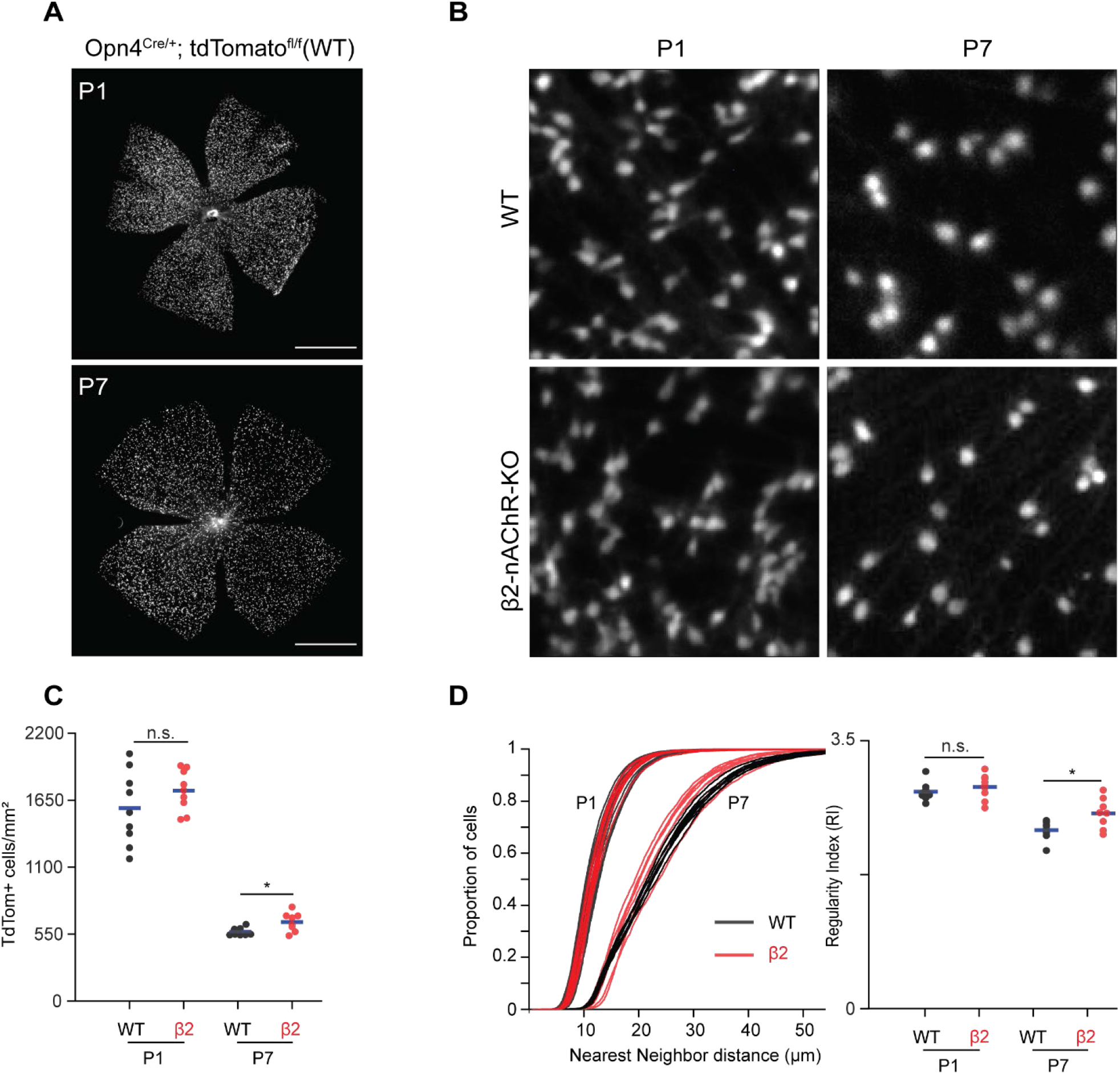
Stage 1 and 2 waves do not contribute to the developmental cell death of ipRGCs. **A**. Example epifluorescence images of Opn4^Cre/+^; tdTomato^fl/fl^ (WT) retinas at P1 and P7. Scalebars are 500 μm; each field of view is 4.7×4.7 mm. **B**. Representative 200×200 μm^2^ fields of view for P1 and P7 WT andß2-nAChR-KO retina. **C**. Average ipRGC densities across different ages and genotypes. p = 0.012. **D**. Cumulative distribution function of nearest neighbor distances (NND) and regularity index (mean NND/SD) from individual retinas, separated by genotype and age. p = 0.020.

We observed a dramatic decrease in the density of ipRGCs in WT retinas from P1 to P7 (Figures 5B, C), consistent with previous studies of ipRGCs (Chen et al., 2013) and which coincides with peak levels of RGC apoptosis, the primary cause of RGC death during development (Young, 1984; Braunger et al., 2014). We found that β2-nAChR-KO retinas exhibited the same density of ipRGCs at P1 as WT retinas, suggesting that the decrease in activity in Stage 1 waves does not regulate ipRGC cell density. At P7, we observed a small but significant increase in ipRGC densities at P7 in β2-nAChR-KO mice than in WT mice (568± 33 ipRGCs/µm^2^ in WT vs 654 ± 78 ipRGCs/µm^2^ in β2-nAChR-KO, n= 8 retinas in each genotype). We cannot determine whether this small difference is due to the smaller size of retinas in β2-nAChR-KO retinas (Xu et al., 1999) or reflects a true increase in cell number. Overall, these data indicate that the cell death processes that regulate ipRGC number during the first postnatal week are not strongly dependent on retinal waves.

To determine the impact of this developmental decrease in cell density on the mosaic organization of ipRGCs, we computed the regularity index, which is equal to the average nearest neighbor distance divided by the standard deviation. A large regularity index is associated with a non-random distribution of somas. Despite the expected increase in nearest neighbor distance in WT and β2-nAChR-KOs retina between P1 and P7 (Figure 5D, see Table 2), there was only a small decrease in the regularity index. Interestingly, the measured mean regularity indices of 2.8 ± 0.12 (WT P1), 2.9 ± 0.16 (β2-nAChR-KO P1), 2.3 ± 0.13 (WT P7), and 2.6 ± 0.2 (β2-nAChR-KO P1) are regular, as they fall within the range of what would be predicted by a random distribution of cells with soma diameters between 7-10 μm (Keeley et al., 2020). Hence the decrease in cell density does not appear make the soma organization more ordered. This might be expected, since our analysis does not differentiate between ipRGC subtypes, each of which likely forms an independent retinal mosaic. Together these data indicate that although retinal waves provide a robust source of depolarization for embryonic and early postnatal ipRGCs, reducing wave activity does not significantly influence the cell death process.

**Table 2:** Nearest-neighbor distances and regularity indices for ipRGCs labeled in Opn4cre::Td-tomato mice.

| | Median nearest neighbor distance ($\mu\text{m}$) | Regularity Index (NND/SD) | Number of retinas |
| --- | --- | --- | --- |
| WT P1 | 11.59± 0.93 | 2.83±0.12 | 9 |
| $\beta 2$ P1 | 11.51±0.55 | 2.89 ±0.16 | 9 |
| WT P7 | 22.32±0.58 | 2.33±0.13 | 8 |
| $\beta 2$ P7 | 20.96±1.40 | 2.55±0.20 | 8 |

Data represented as averages ± SD. This data complements quantification in Figure 5.

## DISCUSSION

We show that Stage 1 retinal waves are a robust source of spontaneous activity in the embryonic retina. Stage 1 waves initiate throughout the retina, propagate over finite regions of varying size and drive periodic depolarizations of neurons in the immature retinal ganglion cell layer. Stage 1 waves are reduced but persist in the presence of gap junction antagonists. Stage 1 waves are blocked by general nAChR antagonists but persist, albeit with reduced cell participation, in a nAChR antagonist that targets α4β2 containing nAChRs. We found that the β2-nAChR-KO mouse, which exhibits strongly reduced Stage 2 waves, also exhibits reduced Stage 1 waves that are insensitive to general nAChR antagonists but are completely blocked by gap junction antagonists, indicating that the electrical synapses may compensate for a lack of at least some nAChRs. Finally, we showed that ipRGCs are depolarized by Stage 1 waves, but that decreasing number of ipRGCs across early postnatal development was unaffected in the β2-nAChR-KO mouse, indicating that the proliferation and cell death processes that influence ipRGC number are not dependent on wave activity.

### Distinctions and similarities between Stage 1 and Stage 2 waves

Our findings support a model in which there are both key differences and some similarities between Stage 1 and 2 waves. The distribution of IEI is peaked at roughly 80 seconds for Stage 1 waves which is a slightly longer interval than the peak interval reported for Stage 2 (compare Figure 1C here to Figure 7D in Ford et al, 2012). Stage 1 and 2 waves also propagate at similar speeds. Key differences are that spontaneous activity between E16 and E18 included small non-propagating events and propagating waves of a broad range of sizes. In contrast, after P1, there are no small non-propagating events, and Stage 2 waves mostly propagate over large areas of the retina, as observed both *in vitro* (Feller et al., 1997; Hilgen et al., 2017) and *in vivo* (Ackman et al., 2012). Note the spatiotemporal properties of Stage 2 waves also vary dramatically across the developmental period spanning P1-P10 (Hilgen et al., 2017; Ge et al., 2021) and therefore we are making these comparisons to the earlier Stage 2 waves.

The similarity in frequency of events and propagation speed indicate that the mechanisms responsible for initiating and propagating large waves are similar for Stage 1 and 2 waves. Indeed, Stage 1 and 2 waves share a dependence on nAChR signaling. Starburst amacrine cells (SACs) are the sole source of ACh in the retina. During the period of development concurrent with Stage 1 waves, SACs begin to migrate toward the INL and send projections to the IPL via homotypic contacts (Ray et al., 2018). Also during this period, SACs start to express choline acetyltransferase at E17 in the rat retina (Kim et al., 2000), equivalent to E15.5 in mice (Schneider and Norton, 1979), and show a response to nicotine in the fetal rabbit retina between E20-27 (Wong, 1995), corresponding to Stage 1 and 2 waves in rabbit (Syed et al., 2004). Together these studies support the idea that during embryonic development, SACs are not only releasing ACh but also forming a cholinergic network, similar to that of Stage 2 wave propagation (Ford and Feller, 2012). Here, we have shown that Stage 1 waves fail to initiate in the presence of the general nAChR antagonists, hexamethonium (Hex) and epibatidine (EPB). This suggests that spontaneous depolarization of SACs is important for wave initiation.

One key difference in ACh signaling between Stage 1 and 2 waves is their sensitivity to the specific nAChR antagonist, DHβE. Though DHβE reduced the number of large propagating Stage 1 waves, many smaller propagating and non-propagating events persisted. In contrast, DHβE is a potent blocker of all activity during Stage 2 waves (Ford et al., 2012). DHβE is an nAChR antagonist with a greater affinity for nAChRs containing α4 and β2 subunits in heterologous systems (Papke et al., 2010; Ho et al., 2020). The fact that all Stage 1 wave activity is blocked by Hex and EPB suggests that different subunit combinations of nAChRs, on both SACs and RGCs, are independently contributing to either the initiation or propagation of Stage 1 waves.

In our hands, global blockade or desensitization of nAChRs completely abolished Stage I waves. This result appears to conflict with previous studies in mice showing that application of curare, a competitive antagonist for nAChRs, preserves small non-propagating events (Bansal, 2000). One possibility is that in contrast to hexamethonium, curare has mixed affinity for neuronal nAChRs. A second is that curare was acting via other neurotransmitter receptors where it has some cross-reactivity not shared by the receptor antagonists that we used (Wotring and Yoon, 1995; Spirova et al., 2019).

In addition to the ACh signaling, gap junctions also play a role in mediating Stage 1 retinal waves. Here we use the gap junction antagonist meclofenamic acid (MFA), which was previously shown to reversibly block junctional conductance (Veruki and Hartveit, 2009), dye coupling (Pan et al., 2007) and spikelets between developing RGCs (Caval-Holme and Feller, 2019). Application of MFA led to a significant reduction in both the frequency and size of Stage 1 waves. Unlike the complete block we observed in Hex and EPB, small propagating and non-propagating waves persisted in MFA, suggesting that gap junctions are important for setting the frequency and area of Stage 1 waves. However, there is the important caveat that MFA can also have some off-target effects that might impact wave propagation (Kuo et al., 2016). The results we observed in MFA are consistent with the pharmacological studies of Stage 1 waves in other species: Stage 1 waves in rabbit are insensitive to nAChR antagonists (Syed et al., 2004), as are early waves in developing chick retina (Catsicas et al., 1998). Note that waves in these species were also sensitive to antagonists of various metabotropic receptors, indicating that neurotransmitters are still important for propagating waves in these systems.

We also observed a difference between Stage 1 and 2 waves in β2-nAChR-KO retinas. Large propagating waves persisted in Stage 1 but they were less frequent and somewhat smaller than those observed in WT. Though the spatiotemporal properties of Stage 2 waves recorded *in vitro* for P1-P8 β2-nAChR-KO retinas is highly dependent on recording conditions with results ranging from sparse activity to high frequency (Bansal et al., 2000; Stafford et al., 2009; Xu et al., 2016), *in vivo* waves in β2-nAChR-KO are infrequent and weakly depolarizing (Burbridge et al., 2014). The activity that persists in β2-nAChR-KO retinas both embryonically and postnatally is resistant to nAChR antagonists, as was observed previously (Bansal et al., 2000). Rather the activity that persists in the β2-nAChR-KO retinas was completely blocked by MFA, therefore suggesting that stage I and II waves in β2-nAChR-KO retinas rely solely on gap junctions (Kirkby and Feller, 2013).

We propose a model for Stage 1 waves consistent with these observations (Figure 6). Stage 1 wave initiation is dependent on the spontaneous depolarization of SACs, but wave propagation is dependent on both gap junction coupling and nAChR activation. Hence Stage 1 wave initiation is similar to Stage 2 waves where the spontaneous depolarization of a SAC activates neighboring SACs, leading to the volumetric release of ACh responsible for wave propagation both in the INL and GCL (Ford et al., 2012). The model for Stage 1 wave propagation is also similar to Stage 2 wave propagation, though a complete description of how gap junction coupling is mediating Stage 2 wave propagation is needed. The pharmacological and genetic block/ablation of gap junctions have yet to reveal a phenotype (Singer et al., 2001; Torborg et al., 2005; Blankenship et al., 2011; Kirkby and Feller, 2013; Caval-Holme and Feller, 2019). Interestingly, pure gap junction mediated waves, such as those we observed in the β2-nAChR KO mouse, are considerably slower than waves in WT retina. A recent computational model of gap junction mediated Stage 1 waves, where waves are initiated by RGCs undergoing rare and random depolarizations that propagate entirely via electrical synapses, argues that the speed of propagation is limited by the slow rate at which the junctional currents charge up the membrane capacitance of neighboring RGCs (Kähne et al., 2019). Hence the faster speed of waves mediated by a combination of nAChRs and gap junctions indicates that diffuse release of ACh leads to faster propagation than electrical synapses alone.

**Figure 6.**
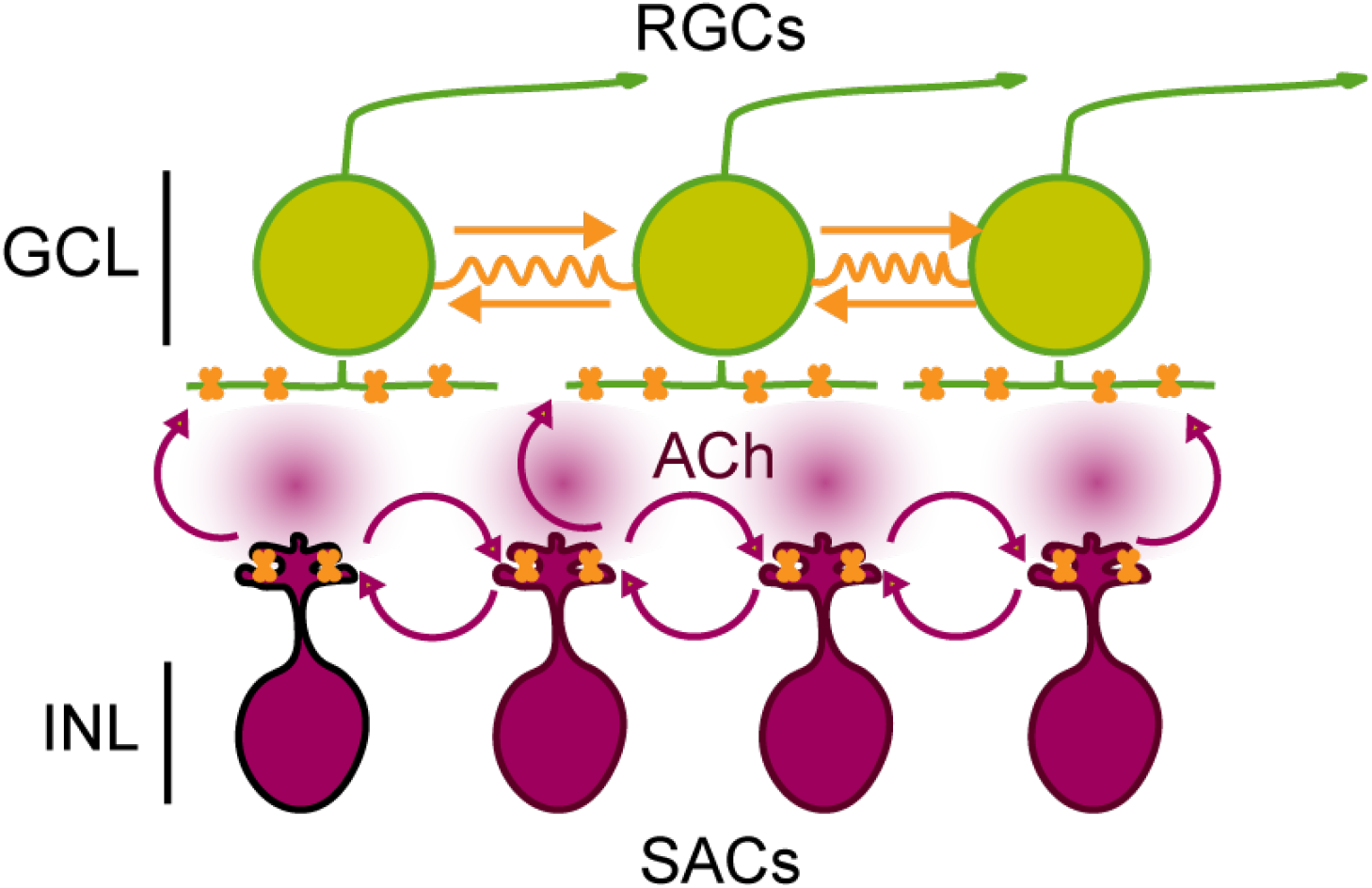
Schematic of Stage 1 wave initiation and propagation. Wave initiation set by spontaneously depolarizing cholinergic amacrine cell/SAC, outlined in black, which release ACh and depolarizes neighboring cells, leading to the volumetric release of ACh. RGCs depolarized by nAChR activation (orange) via SAC-induced ACh release. Wave propagation set by RGC depolarization via gap junctional currents (orange) and volumetric ACh release.

### Interactions between Stage 1 waves and ipRGCs

Early intrinsic light responses of ipRGCs have been implicated in several developmental events (Aranda and Schmidt, 2021), including retinal vascularization (Rao et al., 2013), maturation of circadian circuits (McNeill et al., 2011) and the maturation of the lens to prevent myopia (Chakraborty et al., 2022). Here we report that ipRGCs are robustly activated by Stage 1 waves, similar to our observations that ipRGCs participate in Stage 2 waves (Kirkby and Feller, 2013; Arroyo et al., 2016; Caval-Holme et al., 2022). Thus, it is possible that depolarization via Stage 1 waves may contribute to some of these ipRGC-dependent developmental processes.

To begin to explore to role of Stage 1 and 2 waves in ipRGC development, we monitored the impact of chronically altered waves on the distribution and number of ipRGCs across the retina. Notably, ipRGCs undergo extensive apoptotic cell death, with the peak of apoptosis occurring between P2-P4 (Chen et al., 2013). Prevention of apoptosis during this developmental period doubles the number of ipRGCs and dramatically increases the clumping of M1-ipRGC somas (Chen et al., 2013). In some systems, correlated network activity has been implicated in cell proliferation and cell death. For example, retinal wave activity promotes neurite outgrowth and potentially survival among RGCs (Goldberg et al., 2002). Additionally, in the developing primary somatosensory and motor cortices, higher levels of spontaneous electrical activity were shown to have a neuroprotective effect (Blanquie et al., 2017b). However, here we show in the β2-nAChR-KO mouse, which has significantly diminished Stage 1 and 2 retinal waves, the normal developmental loss of ipRGCs between P1 and P7 is maintained. Thus, Stage 1 and 2 retinal waves are not required for ipRGC apoptosis; however, it is possible that the residual wave activity in β2-nAChR-KO is sufficient to activate pro-survival pathways. A deeper understanding of how spontaneous activity modulates RGC survival pathways (e.g. (Ahmed et al., 2022) is warranted.

## ACKNOWLEDGEMENTS

All authors supported by NIH RO1EY013528, RO1EY019498, P30EY003176. A. T. was supported by K99EY030909. B.E.S. was supported by NIH P30EY003176. We thank members of the Feller Lab for their comments on the manuscript.

## METHODS

### Animals

All animal procedures were approved by the UC Berkeley Institutional Animal Care and Use Committee and conformed to the NIH Guide for the Care and Use of Laboratory Animals, the Public Health Service Policy, and the SFN Policy on the Use of Animals in Neuroscience Research. For our calcium dye based calcium imaging, OPN4cre::tdTomato mice were generated by crossing mice B6.Cg-*Gt(ROSA)26Sor*^*tm9(CAG-tdTomato) Hze*^/J mice from (Jackson Laboratory, Bar Harbor, ME) to the *Opn4*^*Cre/+*^ reporter mouse (T. Schimdt, Northwestern University, Evanston, IL). For our GCaMP6s based calcium imaging, we generated Vglut2::GCaMP6s mice by crossing B6J.129S6(FVB)-*Slc17a6*^*tm2(cre)Lowl*^/MwarJ mice (Jackson Laboratory, Bar Harbor, ME) to B6J.Cg-*Gt (ROSA)26Sor*^*tm96(CAG-GCaMP6s)Hze*^/MwarJ mice (Jackson Laboratory, Bar Harbor, ME).

ipRGC density measurements were conducted on P1 – P7 mice of either sex using OPN4cre::tdTomato mice. To obtain mice that were precisely at the correct embryonic age, we set up timed pregnancies and checked vaginal plugs every morning for four days after the animals were paired. We used the β2-nAChR-KO mouse line in which the β subunit of the nicotinic acetylcholine receptor is knocked out as a genetic model in which cholinergic retinal spontaneous activity is disrupted. For experiments regarding the influence of spontaneous retinal activity on the distribution of ipRGCs across the retina, we used β2^-/-^::OPN4cre::tdTomato (β2-nAChR-KO) mice, generated by crossing β2^-/-^ (A. Beaudet, Baylor University, Waco, TX) mice to OPN4cre::tdTomato mice to label all melanopsin-expressing cells. All mouse lines are maintained on a C57BL/6 genetic background. All animals used for two-photon calcium imaging experiments and immunohistochemistry were housed in 12-hour day/night cycle rooms.

### Retinal preparation

On the day of the experiment, pregnant dams were deeply anesthetized via isoflurane inhalation and fetuses were harvested via a cesarean section. tdTomato positive fetuses were identified using miner goggles (Biological Laboratory Equipment Services and Maintenance Ltd., Model: GFsP-5). Fetuses were kept alive in 50 mL falcon tubes filled with oxygenated (95% O_2_ 5% CO_2_) ACSF (in mM, 119 NaCl, 2.5 KCl, 1.3 MgCl_2_, 1 K_2_HPO_4_, 26.2 NaHCO_3_, 11 D-glucose, and 2.5 CaCl_2_). Fetuses were then euthanized sequentially by decapitation. Eyes were immediately enucleated and retinas were dissected at room-temperature in oxygenated ACSF, under a dissecting microscope. Isolated retinas were mounted whole over a 1–2 mm^2^ hole in nitrocellulose filter paper (Millipore) with the photoreceptor layer side down and transferred to the recording chamber of a two-photon microscope for imaging. The whole-mount retinas were continuously perfused (3 ml/min) with oxygenated ACSF warmed to 32-34°C by a regulated inline heater (TC-344B, Warner Instruments) for the duration of the experiment. Additional retina pieces were kept in the dark at room temperature in ACSF bubbled with 95% O2, 5%CO2 until use (maximum 8 h).

For the calcium imaging experiments, retinas were bath loaded with the calcium indicator Cal 520 AM (AAT Bioquest) for 1-2 hours at 32°C.

### Two-photon calcium imaging

Two-photon fluorescence measurements were obtained with a modified movable objective microscope (MOM) (Sutter instruments, Novator, CA) and made using an Olympus 60X, 1.00 NA, LUMPlanFLN objective (Olympus America, Melville, NY) for single cell resolution imaging (FOV: 203 × 203 um) or a Nikon 16X, 0.80 NA, N16XLWD-PF objective (Nikon, Tokyo, Japan) for large FOV (850 × 850 um) imaging. Two-photon excitation was evoked with an ultrafast pulsed laser (Chameleon Ultra II; Coherent) tuned to 920 nm to image Cal520, GCaMP6s, and tdTomato. Laser power was set between 6.5 mW-12mW for imaging of Cal520 and tdTomato expression. The microscope system was controlled by the ScanImage software (https://www.scanimage.org/). Scan parameters were [pixels/line x lines/frame (frame rate in Hz)]: [256 × 256 (1.48Hz)], at 2 ms/line. This MOM was equipped with a through-the-objective light stimulation and two detection channels for fluorescence imaging.

### Epifluorescent macroscope calcium imaging

Epifluorescent calcium imaging were obtained on a custom built macroscope with an Olympus XLFLUOR4X/340 4X 0.28 NA objective, a Teledyne Kinetix camera. Collectively, this macroscope has 4.7 mm x 4.7 mm FOV, and 1.5 µm/pixel. All movies were taken at a 1 Hz frequency and pixels were binned 4 × 4 bringing the resolution down to 5.9 µm/pixel, still maintaining single cell resolution. Cal520 and GCaMP6s excitation was evoked with a 474 nm LED. A full description and building instructions can be found at: https://github.com/Llamero/DIY_Epifluorescence_Macroscope

### Initiation site measurements

Macroscope recordings of Stage 1 waves were used for the manual detection of initiation sites. Small non-propagating events were identified as local regions of correlated calcium activity with fixed areas and no wave fronts. Small propagating waves were identified as regions of correlated calcium activity with no fixed areas and with wave fronts covering up to 25% of the retina. Large propagating waves were identified as regions of correlated calcium activity with no fixed areas and with wave fronts covering up to 90% of the retina.

### Pharmacology

We blocked gap junctions via application of the gap junction blocker Meclofenamic acid (MFA, 50 µM, Sigma Aldrich). We blocked the nicotinic acetylcholine signaling pathway via application of the broad nicotinic receptor antagonists hexamethonium (Hex, 100 µM, Sigma Aldrich) and epibatidine (EPB, 10 nM, Sigma Aldrich) as well as the specific antagonist dihydro-ß-erythroidine hydrobromide (DHßE, 8 µM, Sigma Aldrich).

The following procedure was used for all pharmacology experiments: We recorded baseline activity in ACSF for 8 minutes before pharmacological agents were applied to the perfusion system. We then waited 15-30 minutes for the agents to take effect before acquiring another 8-minute recording session.

### Image analysis of population calcium imaging movies

Movies were preprocessed for motion correction using a MATLAB code from the Flat Iron Institute (https://github.com/flatironinstitute/NoRMCorre). The baseline movie frame (F0) was computed by taking the temporal median projection of all the movie frames. Each movie frame (F) was normalized by dividing its difference from the baseline frame (F-F0) by the baseline frame ((F-F0)/F0) to produce a ΔF/F0 movie. For movies taken on the two-photon microscope circular ROIs were drawn on all cells within the field of view (FOV). Additional circular ROIs were drawn for tdTomato+ cells. For movies taken on the macroscope a grid of 10 µm x 10 µm squares, that were spaced 1.5 pixels apart, were drawn over the whole surface of the retina using a custom FIJI macro. The ROIs and the ΔF/F0 movie were then imported into MATLAB for further analysis using custom algorithms. Traces for each FOV and ROI were computed as the mean value of the pixels enclosed by the ROI in each frame of the ΔF/F0 movie. A threshold of 10% ΔF/F0 was used to detect waves in each FOV. To determine the significance of the effects of DHßE, Hex, EPB and MFA on wave frequency a Shapiro-Wilk test of normality was performed followed by a student paired t-test. To determine significance of the differential effects of DHßE, Hex and EPB on wave frequency a Shapiro-Wilk test of normality was performed followed by an ANOVA, then a Tukey HSD post hoc test.

To determine if the distributions of cell participation and mean event amplitude for each wave in control and DHßE conditions were significantly different a Shapiro-Wilk test of normality was performed and followed by a two sample Kolmogorov-Smirnov test.

To determine if the mean response amplitude of a cell during a wave and the percentage of waves each cell participated in across retinal ganglion cell populations, we first determined ΔF/F0 values that corresponded to waves by comparing them against a threshold set by the 95^th^ percentile of a bootstrapped baseline. This 95^th^ percentile threshold was computed iteratively for each cell across every FOV. We then calculated the percentage of waves each cell participated in and averaged this value for every FOV. Similarly, mean response amplitudes were calculated for each cell and then averaged for each FOV.

### Analysis of ipRGC densities

To image the number of ipRGCs in fixed retinas, dissected retinas from P1 and P7 mice were fixed in 4% PFA for 30 minutes. The fixed retinas were subsequently mounted on a slide with vectashield and a cover slip, then imaged within an hour of mounting on the macroscope. For P1 retinas, Z-stacks were acquired by manually turning the focus knob.

We first identified the centroid of each ipRGCs. For P7 retinas, where there is more space between cells, we employed the following automatic segmentation. Images were bandpass filtered and somata automatically segmented using the Morpholibj (Legland et al., 2016)classic watershed tool to obtain 8-bit binarized masks. The masks were then processed in MATLAB in order to obtain the centroid locations and nearest neighbor distances for each soma. For P1 retina, where there is less space between cells and in fact cells seem to form clusters, automatic segmentation was not possible. Therefore, cells were manually marked using the ImageJ multipoint tool and soma locations exported as a CSV file. For all ages, the centroid data was imported to MATLAB for further analyses.

Density was quantified by dividing the microscope field of view up into 200×200 μm squares, manually excluding ones that did not cover the retina or covered partial or damaged parts of the retinas. Out of the resulting squares (∼150 per P7 retina, ∼100 per P1 retinas) 75 squares (for P7 retinas) and 50 squares (for P1 retinas) were randomly selected and the average density of TdTom+ cells in those squares calculated.

To quantify the nearest neighbor distances, we used a custom-written MATLAB code that, for each ipRGCs, identified the closest neighbor using the shortest Euclidean distance.

### Statistical Tests

Details of statistical tests, number of replicates, and p values are indicated in the figures and figure captions. P values less than 0.05 were considered significant.

### Supplemental Materials

**Figure 1-figure supplement 1.**
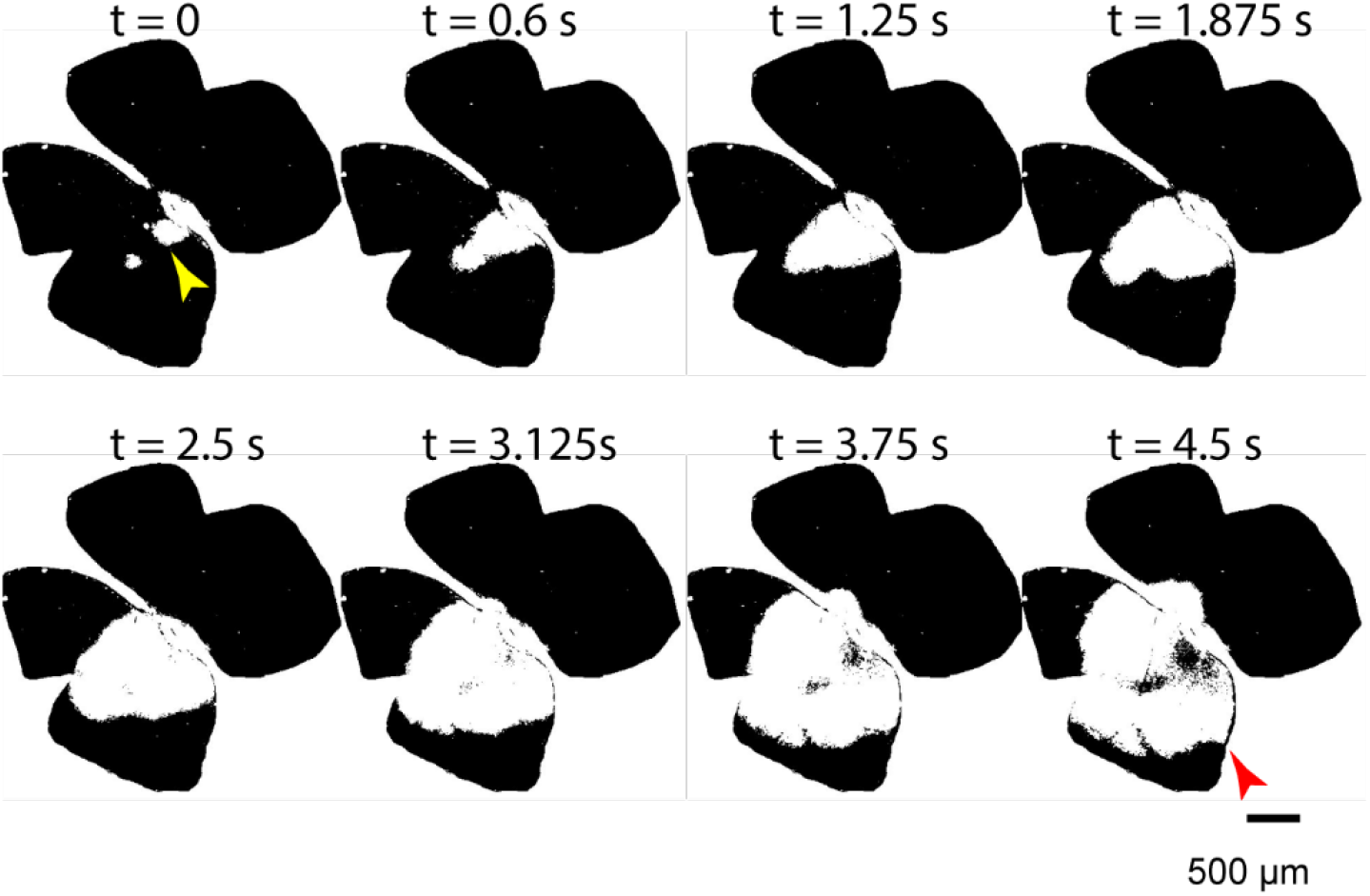
Average wave speed measurement. Temporal progression of a large Stage 1 wave in WT control conditions. The average wave speed was calculated by measuring the distance between two points: the first point was marked at a random location on the edge of area defined early in the wave event (yellow arrowhead; initiation spot); the second point was located at the wave edge that was parallel to the direction of first point on the initiation spot (red arrowhead). The speed was then calculated by dividing this distance by the propagation time of the wave.

**Figure 1-figure supplement 2.**
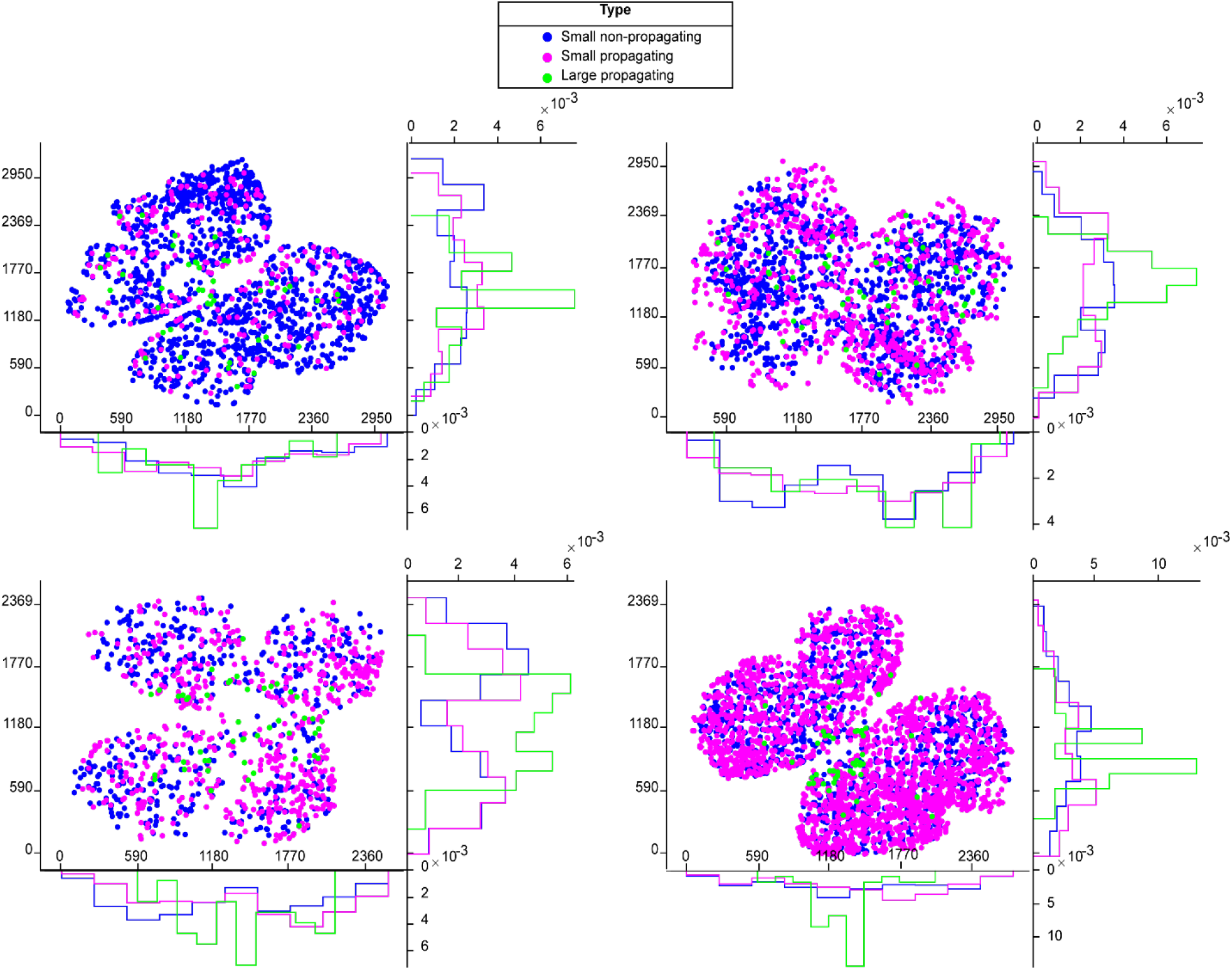
Distribution of Stage 1 wave initiation sites. Scatter plots and histogram showing the distribution of initiation sites for small non-propagating waves (blue), small propagating waves (magenta), and large propagating waves (green) across four E18 Vglut2cre::GCaMP6s retinas. X-axes for the scatter plots and histograms = μm. Y-axes for scatter plots = μm. Y-axes for histograms = proportion of initiation sites.

**Figure 3-figure supplement 1.**
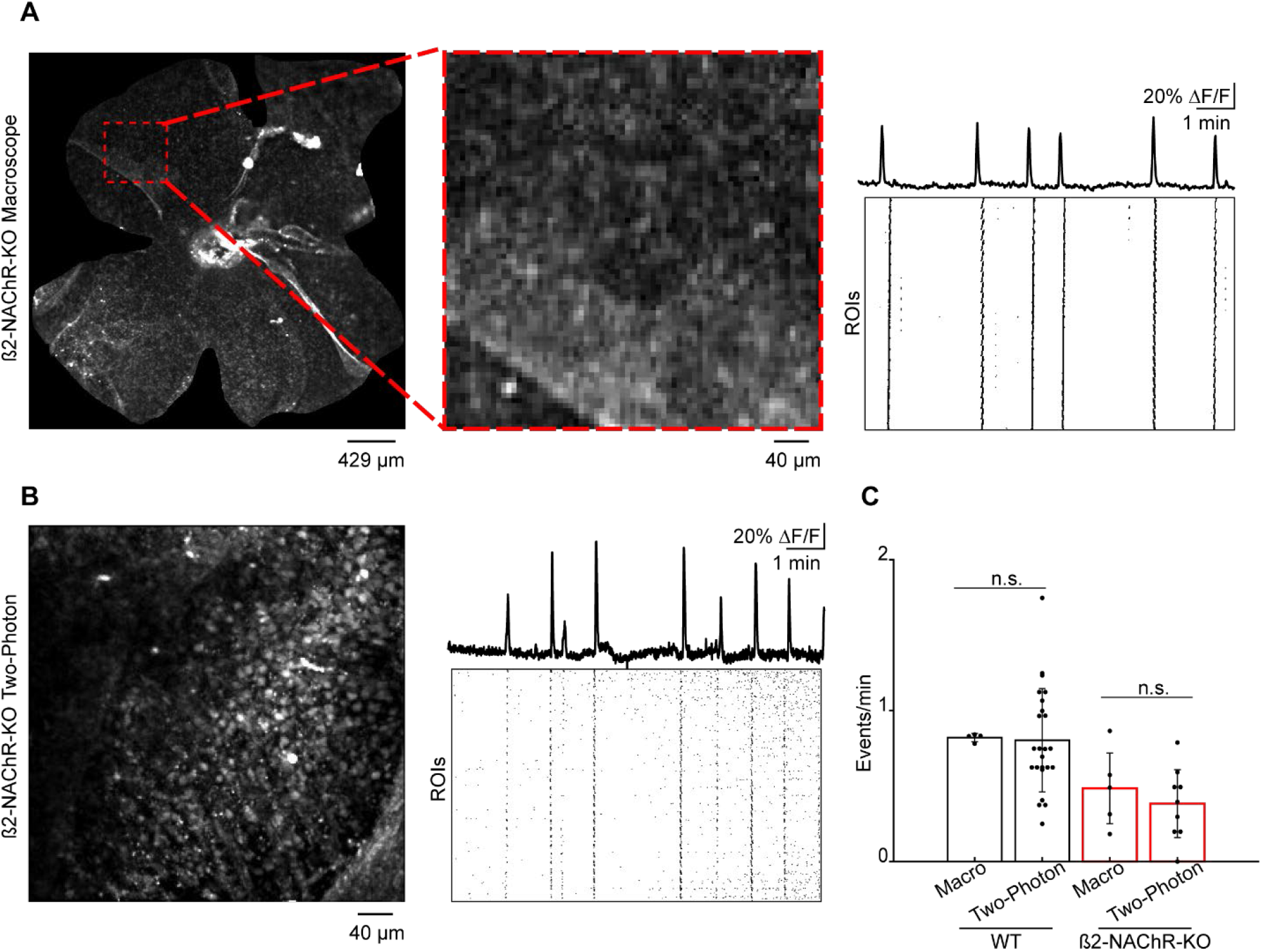
Frequency of Stage 1 waves in WT and β2-nAChR-KO retinas recorded on the macroscope vs the two-photon microscope. **A**. *Left:* example FOV, whole mount β2-nAChR-KO, on the macroscope. Red dashed square = 425 μm^2^ FOV, equivalent to two-photon FOV. *Middle:* magnified FOV. *Right:* normalized fluorescent traces showing Ca^2+^/wave activity and raster plot summarizing Ca^2+^/wave activity across 864 square ROIs (10 μm x 10 μm) on the macroscope. **B**. *Left:* example FOV, whole mount β2-nAChR-KO, on the two-photon microscope. *Right:* normalized fluorescent traces showing Ca^2+^/wave activity and raster plot summarizing Ca^2+^/wave activity across 864 square ROIs (10 μm x 10 μm) on the two-photon microscope. **C**. Bar graph showing frequency of waves/events on the macroscope and two-photon microscope in WT and β2-nAChR-KO. WT: n = 4 retinas (macroscope; Average ± SD: 0.82 ± 0.03), n = 24 retinas (two-photon; Average ± SD: 0.80 ± 0.34), p = 0.92. β2-nAChR-KO: n = 5 retinas (macroscope; Average ± SD: 0.48 ± 0.23), n = 9 retinas (two-photon; Average ± SD: 0.38 ± 0.22), p = 0.48.

**Figure 5-figure supplement 1.**
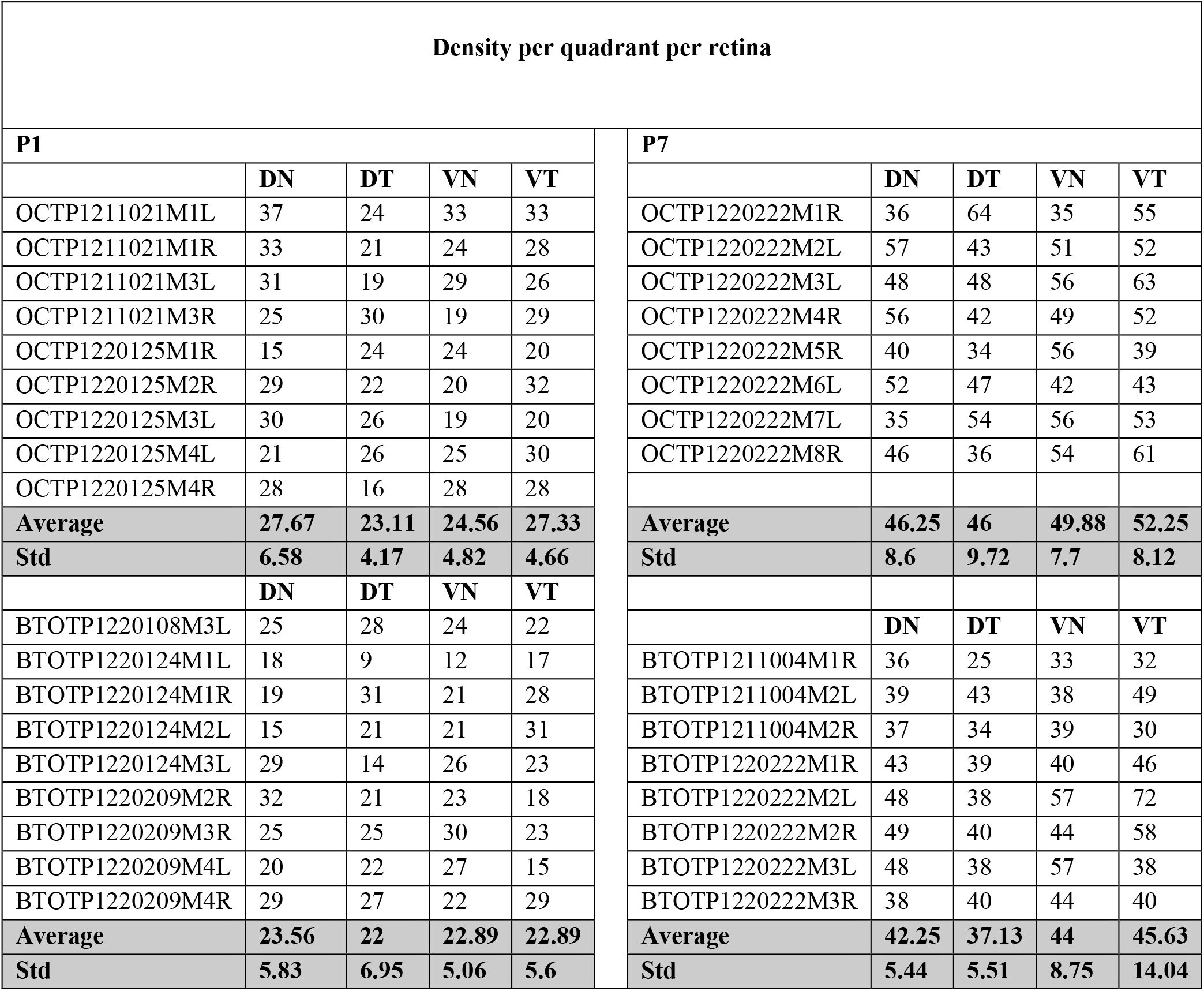
Density of tdTom+ cells per quadrant. Density analysis of per retinal quadrant shows no significant difference in density across quadrants.

